# Simultaneous carbon catabolite repression governs sugar and aromatic co-utilization in *Pseudomonas putida* M2

**DOI:** 10.1101/2023.05.23.541960

**Authors:** Shilva Shrestha, Deepika Awasthi, Yan Chen, Jennifer Gin, Christopher J. Petzold, Paul D. Adams, Blake A. Simmons, Steven W. Singer

## Abstract

*Pseudomonas putida* have emerged as promising biocatalysts for the conversion of sugars and aromatics obtained from lignocellulosic biomass. Understanding the role of carbon catabolite repression (CCR) in these strains is critical to optimize biomass conversion to fuels and chemicals. The CCR functioning in *P. putida* M2, a strain capable of consuming both hexose and pentose sugars as well as aromatics, was investigated by cultivation experiments, proteomics, and CRISPRi-based gene repression. Strain M2 co-utilized sugars and aromatics simultaneously; however, during co-cultivation with glucose and phenylpropanoid aromatics (*p-*coumarate and ferulate), intermediates (4-hydroxybenzoate and vanillate) accumulated, and substrate consumption was incomplete. In contrast, xylose-aromatic consumption resulted in transient intermediate accumulation and complete aromatic consumption, while xylose was incompletely consumed. Proteomics analysis revealed that glucose exerted stronger repression than xylose on the aromatic catabolic proteins. Key glucose (Eda) and xylose (XylX) catabolic proteins were also identified at lower abundance during co-cultivation with aromatics implying simultaneous catabolite repression by sugars and aromatics. Downregulation of *crc* via CRISPRi led to faster growth and uptake of glucose and *p*-coumarate in the CRISPRi strains compared to the control while no difference was observed on xylose + *p*-coumarate. The increased abundance of the Eda and amino acids biosynthesis proteins in the CRISPRi strain further supported these observations. Lastly, small RNAs (sRNAs) sequencing results showed that CrcY and CrcZ homologues levels in M2, previously identified in *P. putida* strains, were lower under strong CCR (glucose + *p*-coumarate) condition compared to when repression was absent (*p*-coumarate or glucose only).

**IMPORTANCE:** A newly isolated *Pseudomonas putida* strain, *P. putida* M2, can utilize both hexose and pentose sugars as well as aromatics making it a promising host for the valorization of lignocellulosic biomass. Pseudomonads have developed a regulatory strategy, carbon catabolite repression, to control the assimilation of carbon sources in the environment. Carbon catabolite repression may impede the simultaneous and complete metabolism of sugars and aromatics present in lignocellulosic biomass and hinder the development of an efficient industrial biocatalyst. This study provides insight into the cellular physiology and proteome during mixed-substrate utilization in *P. putida* M2. The phenotypic and proteomics results demonstrated simultaneous catabolite repression in the sugar-aromatic mixtures while the CRISPRi and sRNA sequencing demonstrated the potential role of the *crc* gene and small RNAs in carbon catabolite repression.

## 1. INTRODUCTION

Lignocellulosic biomass, including residue from agriculture and forestry, is an attractive alternative carbon feedstock for bioprocesses (1). However, its inherent heterogeneity and recalcitrant nature pose a unique challenge to efficient utilization of this abundant, renewable resource (2). Integrating physicochemical pretreatment with microbial conversion of lignocellulose-derived sugars and aromatics to fuels and chemicals has been proposed to overcome this challenge (3, 4). Depolymerization of lignocellulose results in mixtures of hexose (glucose) and pentose (xylose, arabinose) sugars as well as aromatics (4). Most microbes grow on a subset of these compounds which limits their industrial use. *Pseudomonas alloputida* KT2440 (previously known as *Pseudomonas putida* KT2440), hereafter referred to as KT2440, is a promising industrial biocatalyst due to its metabolic versatility, ability to tolerate environmental stresses, and amenability to genetic manipulation (5–8). However, KT2440 cannot natively utilize xylose (9), which makes up a significant fraction of total plant sugars, preventing the complete utilization of lignocellulose sugars. Recently *P. putida* strains capable of growing on pentose sugars were characterized by isolation, genomic sequencing, and phylogenetic analysis (10). One of these strains, *Pseudomonas putida* M2 (hereafter referred to as M2), is a newly isolated soil bacteria that grows on glucose, xylose, and arabinose as well as biomass-derived aromatic monomers, making it a promising host for plant biomass valorization (10, 11).

To thrive in different nutritional environments with numerous and fluctuating amounts of carbon sources, Pseudomonads regulate carbon substrate use to optimize metabolism and growth. This complex regulation is known as carbon catabolite repression (CCR) (12, 13). Previous studies have shown that Pseudomonads generally prefer organic acids and amino acids to glucose, which is preferred to hydrocarbons and aromatic compounds, with a few exceptions (12, 14–16). For instance, KT2440 utilizes glucose and acetate in preference to aromatics (14, 17, 18). CCR also influences other behavior such as virulence, antibiotic resistance, quorum sensing, stress response, bioremediation pathway, and intracellular communication (13, 19). In Pseudomonads, Crc (catabolite repression control) is the major regulatory protein that inhibits the translation of mRNAs responsible for the catabolism of non-preferred compounds in presence of other preferred carbon sources (12). The RNA-binding Hfq protein recognizes the catabolite activity (CA) motif-containing AANAANAA in the target mRNA thus facilitating a Crc-dependent inhibition of mRNA translation (20, 21). The presence of small RNAs (sRNAs) that contain several CA motifs antagonizes the regulatory effect of Crc by sequestering Hfq and thus determine the strength of Crc-mediated CCR (22).

Depending on the type of lignocellulosic biomass and depolymerization method employed, a heterogeneous mixture of sugar and aromatic monomers is obtained. The conversion of these mixtures makes understanding cellular physiology during mixed-substrate utilization of particular interest. To enable the conversion of lignocellulose-derived sugars and aromatics with a single microbe, the microbial host must consume all the carbon sources present rapidly and simultaneously. However, preferential or incomplete consumption of carbon sources due to CCR hinders efficient valorization of lignocellulosic biomass-derived components (14, 23) and negatively affects the economics of the bioconversion process. Accumulation of intermediates due to CCR creates challenges in the downstream processes used to extract the desired product. A systemic understanding of CCR in M2 is required to overcome this barrier and develop strategies for metabolic engineering to develop a robust microbial host. This work focused on characterizing and comparing the growth of M2 when both biomass-derived sugar and aromatic compounds were simultaneously present in the growth medium. Furthermore, CRISPRi and small RNA (sRNA) sequencing were employed to probe the role of *crc* and sRNAs in CCR-mediated repression in M2.

## 2. RESULTS

### Sugars and aromatics were simultaneously utilized in *P. putida* M2

Strain M2 grows on both hexose (glucose) and pentose sugars (xylose, and arabinose) (10) and biomass-derived aromatics monomers (*p*-coumarate (*p*CA), 4-hydroxybenzoate (4-HBA), ferulate (FA), and vanillate (VA)) (Figure S1), consistent with metabolic pathways identified in its genome (10). Previous studies have shown that Pseudomonads such as KT2440 prefer sugars over aromatics (14, 17, 18), which complicates the development of an industrially-relevant strain capable of efficiently utilizing both sugar and aromatics. We, therefore, investigated the growth phenotypes of M2 in a mixture of glucose/xylose + monoaromatics.

Both glucose and aromatics (*p*CA, 4-HBA, FA, or VA) were simultaneously metabolized (Figures 1 and S2) in M2; however, both compounds were not consumed completely after 48 h. On average, 52.1-76.9% of glucose was depleted while 58.6-81.7% of aromatics was consumed over a period of 60 h. Also, M2 exhibited a longer lag phase in glucose and aromatic mixtures compared to growth in single substrate conditions (Figures 1, S1, and S2). M2 exhibited a 24 h lag time when grown in glucose + *p*CA after which simultaneous consumption of both substrates commenced. In contrast, growth on xylose and aromatics showed no lag, and the aromatics were consumed faster than the xylose. After 48h, xylose consumption was not complete, as was observed for glucose. For example, 100% of the *p*CA was depleted within 9 h in growth experiments with xylose + *p*CA, at which time approximately 29% of xylose was consumed.

**Figure 1.**
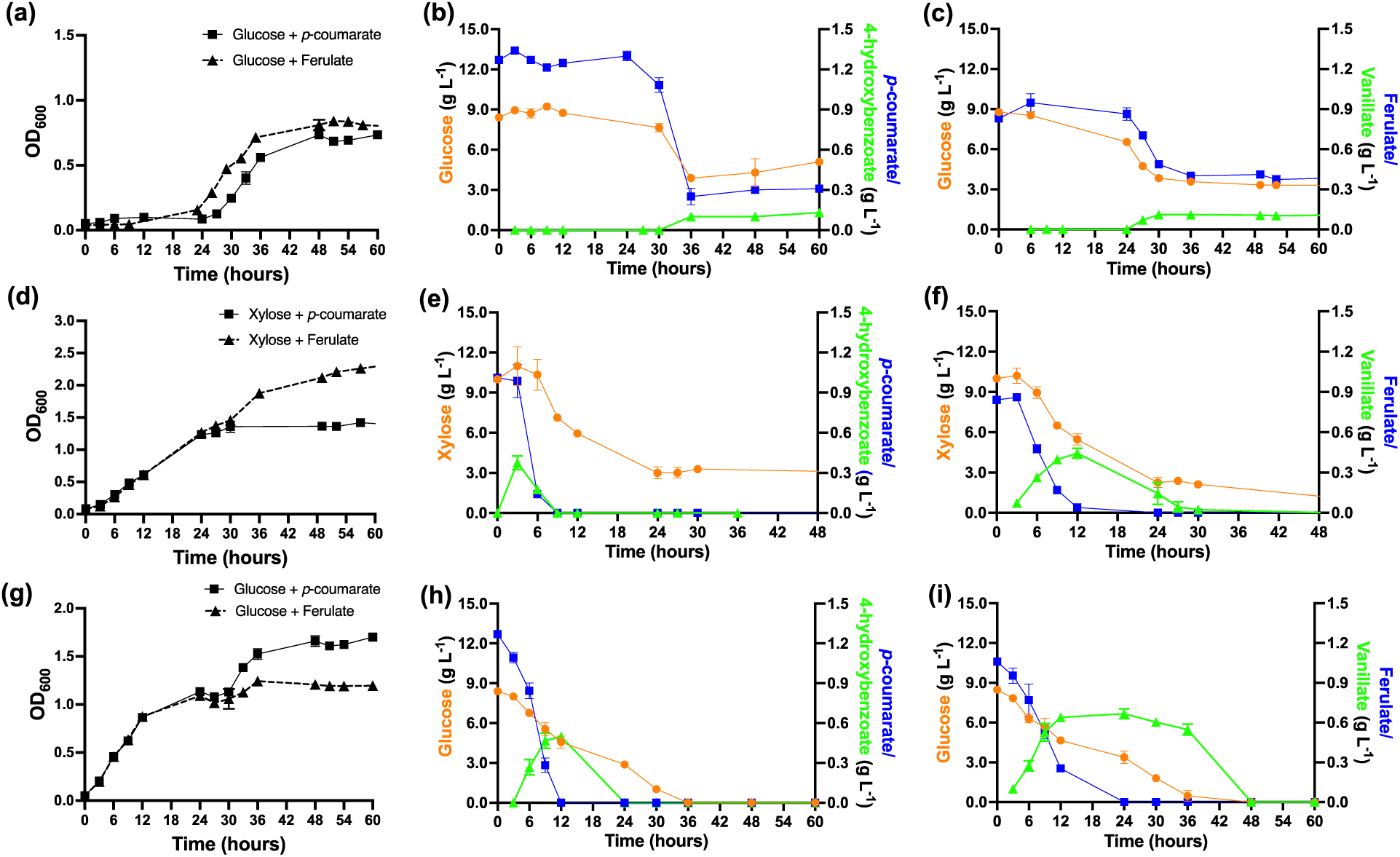
Phenotype of *P. putida* M2 (a-f) and *Pseudomonas alloputida* KT2440 (g-i) showing OD_600_ and substrate consumption and accumulation of intermediates in a mixture of Glucose + *p*-coumarate (b, h), Glucose + Ferulate (c, i), Xylose + *p*-coumarate (e) Xylose + Ferulate (f) as carbon sources. The data represent the mean value of three replicates and error bars represent the standard deviation.

During growth on both glucose and xylose, aromatic intermediates 4-HBA (in *p*CA) and VA (in FA), accumulated during the cultivation. In the case of glucose, the aromatic intermediates persisted as the uptake of *p*CA and FA ceased (Figures 1b and 1c). For xylose growth, transient accumulation of 4-HBA and VA was observed (Figures 1e and 1f). The co-consumption of glucose and aromatics in strain M2 was compared to co-consumption by KT2440 under the same conditions. Both glucose and aromatics were consumed simultaneously in KT2440 with aromatics consumed at a faster rate; unlike M2, both substrates were completely metabolized (Figures 1h and 1i). Furthermore, the metabolic intermediates 4-HBA and VA accumulated transiently in KT2440 during the growth consistent with previous studies (14, 17).

### Aromatic and sugar catabolic proteins were present at significantly lower abundance during their co-utilization

Phenotypic growth data of M2 and KT2440 on sugars and aromatics indicated simultaneous CCR was occurring. To study these effects on protein levels, the proteome of M2 and KT2440 grown in dual carbon conditions (sugar + aromatics, Table S1) were analyzed and compared with those grown in minimal media containing sugar only or aromatics only (control). We assessed the differentially abundant proteins using a criterion of p ≤ 0.05 and log_2_ fold change (FC) ≥ 0.5 value (see Supplementary Information Tables S2-S21 for a complete list of the differentially abundant proteins).

For M2, the proteins involved in the catabolism of *p*CA and FA to 4-HBA and VA, respectively, such as feruloyl-CoA synthase (Fcs), feruloyl-CoA hydratase-lyase (Ech), and vanillin dehydrogenase (Vdh) were present at lower abundance (−1.3 to −3.1 log_2_ FC) in glucose + *p*CA and glucose + FA conditions compared to the control cells grown in *p*CA or FA only (Figures 2a and 2b, Tables S2 and S4). The abundances of 4-hydroxybenzoate hydroxylase (PobA) and vanillate O-demethylase monooxygenase subunit (VanA) responsible for converting 4-HBA and FA, respectively, to protocatechuate (PCA), were also significantly lower (−1.9 log_2_ FC for PobA and −2.3 log_2_ FC for VanA). These results are in agreement with the accumulation of intermediates 4-HBA and VA observed during the growth in glucose + *p*CA or FA mixture (Figure 1). Many of the proteins in the PCA branch of the β-ketoadipate pathway (PcaHG, PcaB, PcaC, and PcaD) and β-ketoadipate conversion to the tricarboxylic acid (TCA) intermediates (PcaIJ and PcaF) were also found at significantly lower abundances in dual substrate conditions compared to the control even though PCA was not observed to accumulate during growth. Similar results were observed for M2 cells grown on glucose + 4-HBA or VA where PobA (−1.7 log_2_ FC), Van A (−2.8 log_2_ FC), and most of the β-ketoadipate pathway proteins (Figure S3, Tables S3 and S5) were significantly at lower abundance in dual substrate growth. For KT2440, the aromatic catabolic proteins (−1.0 to −2.5 log_2_ FC) and most of the β-ketoadipate (−0.6 to −2.4 log_2_ FC) proteins, with the notable exception of Ech, which is involved in the catabolism of both *p*CA and ferulate were also present at significantly lower abundance (Figure S4, Tables S18-S21). The lower abundances observed during glucose/aromatic growth in KT2440 were consistent with previous proteomic studies of substrate co-utilization (14, 17).

**Figure 2.**
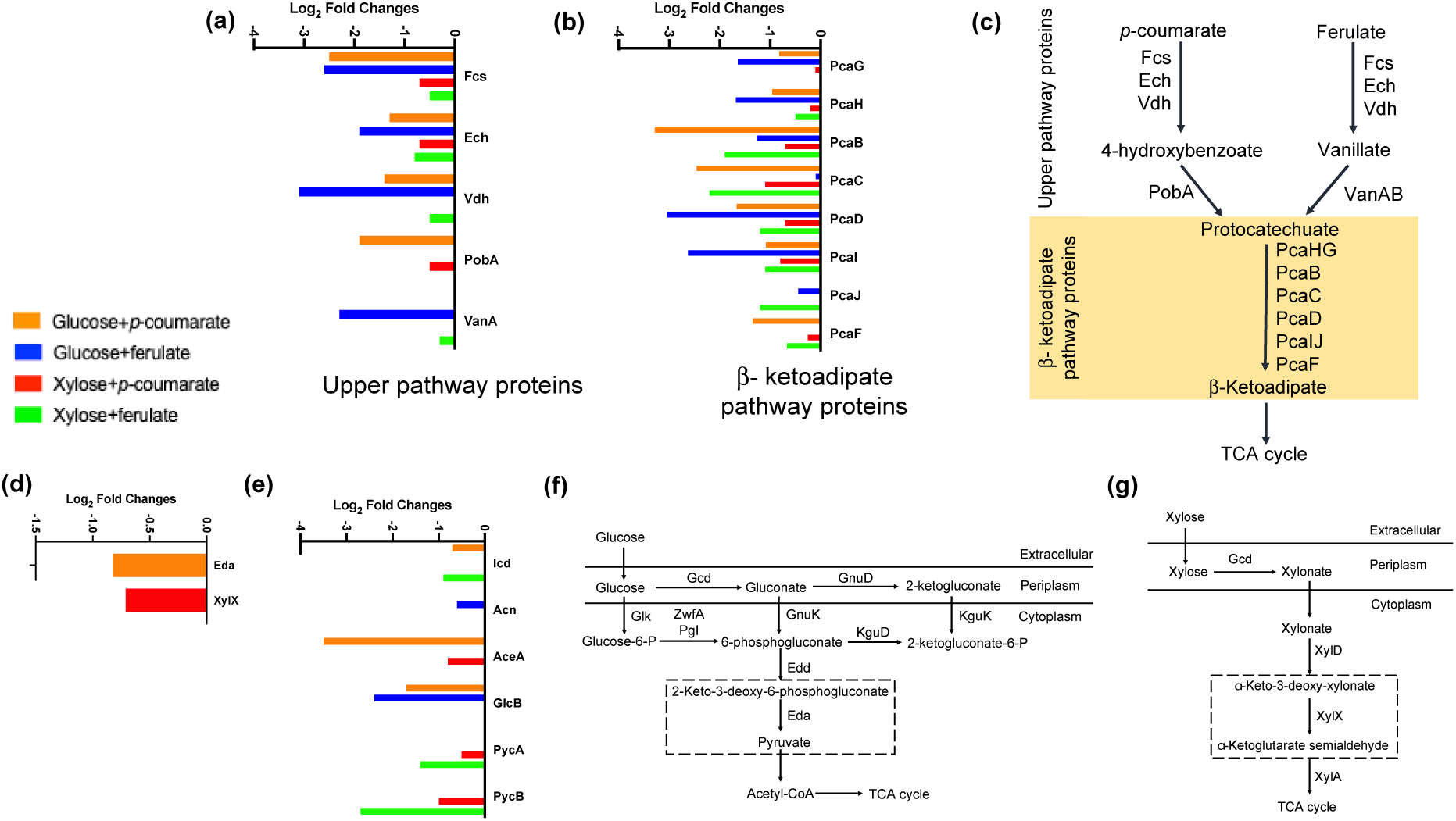
The significantly (p ≤ 0.05, log_2_FC ≤ 0.5) lower abundance upper pathway proteins (a) and β-ketoadipate pathways proteins (b) involved in aromatic catabolism, glucose/xylose catabolic proteins (d), and tricarboxylic acid cycle and gluconeogenesis proteins (e) in *Pseudomonas putida* M2 grown in dual carbon source condition when compared to single carbon source. The schematics of aromatic (c), glucose (f), and xylose (g) catabolic pathways in *P. putida* M2 are shown as described previously (10). The log_2_FC was calculated from three biological replicates for each condition.

Comparing the proteome of M2 cells grown in glucose + aromatics with glucose as the sole carbon source identified that 2-keto-3-deoxy-6-phosphogluconate (KDPG) aldolase (Eda), which converts KDPG into glyceraldehyde-3-phosphate and pyruvate, was present at significantly lower abundance(−0.8 log_2_FC, Figure 2d, Table S6), consistent with incomplete glucose consumption. A subset of TCA and glyoxylate cycle proteins (isocitrate dehydrogenase (Icd), aconitate hydratase (Acn), malate synthase (GlcB), and isocitrate lyase (AceA)) were also observed in significantly lower abundance compared to the single carbon conditions (Figure 2e). Additionally, some amino acid biosynthesis proteins (2-isopropylmalate synthase, glutamine synthetase, O-succinylhomoserine sulfhydrylase), catabolic proteins (ornithine decarboxylase and ornithine carbamoyltransferase), and transporters were also significantly at lower abundance (Tables S2-S9).

When grown in minimal media containing xylose and *p*CA or FA, only some of the aromatic catabolic and β-ketoadipate proteins were present at lower abundance, unlike what was observed in the presence of glucose (Tables S10-S13, Figures 2a and 2b). Interestingly, pyruvate carboxylase subunits A and B proteins, which catalyze the first step of gluconeogenesis and replenish TCA cycle intermediates, were also found at significantly lower abundances (−0.5 to −1.9 log_2_ FC for subunit A and −1.0 to −2.7 log_2_ FC for subunit B) (Figure 2e). For xylose metabolism, co-cultivation with *p*CA lowered the relative abundance of 2-keto-3-deoxyxylonate dehydratase (XylX) (−0.7 log_2_FC, Figure 2d, Table S14), a key xylose catabolic protein which dehydrates α-keto-3-deoxy-xylonate to α-ketoglutarate semialdehyde, thus explaining the incomplete consumption of xylose. Amino acid biosynthesis (threonine synthase, 2-isopropylmalate synthase, acetylornithine/N-succinyldiaminopimelate aminotransferase, glutamate N-acetyltransferase/amino-acid N-acetyltransferase, tryptophan synthase alpha chain) and transport proteins, as well as ribosomal proteins, were present at significantly lower abundance during xylose + aromatics growth compared to when only single carbon was present.

### *crc* downregulation via CRISPRi enhanced growth and metabolism in dual carbon substrates

As Crc has been identified as the main regulatory protein governing CCR in Pseudomonads (12), the *crc* gene was downregulated using a previously developed dual-inducible CRISPR/dCas9-based gene repression system to determine if the aromatic and sugar metabolism in M2 are subject to Crc regulation (24, 25). The functionality of the arabinose-inducible pRGPspdCas9bad-edd plasmid (the backbone plasmid used to design CRISPRi strains) was confirmed to verify sgRNA expression (Figure S5). The pRGPspdCas9bad-edd strain showed delayed growth compared to the control due to sgRNA targeted repression of the *edd* gene, an essential gene for growth on glucose.

We compared the growth phenotype (Figures 3 and S6) and proteomes (Table S22) of the CRISPRi strains to the control strain containing empty plasmid (pRGPspdCas9bad, Table S23) in a minimal medium containing glucose + *p*CA or FA and xylose + *p*CA. The CRISPRi strains (crc-1, crc-2, and crc-3) grew faster than the control strain upon induction when fed with a 10:1 mixture of glucose + *p*CA or FA. The CRISPRi strains grew within 9-12 h while the control strain exhibited a much longer lag phase (32-36 h, Figures 3 and S6). However, the growth profiles were similar between the CRISPRi and control strains in the presence of xylose + *p*CA. The substrate consumption results also showed that both glucose and aromatics (*p*CA or FA) were consumed faster by the CRISPRi strains than the control while xylose and *p*CA consumption profiles were identical in all strains, thus corroborating the OD_600_ measurements. However, both glucose and *p*CA/FA were still not completely consumed in both the CRISPRi strains as observed in the control experiment. It should be noted that the expression of the sgRNA targeting *crc* gene was under the control of the arabinose inducible P*_araC_/_bad_* promoter system. Even though M2 can use arabinose as a carbon source for its growth, we confirmed that arabinose was not consumed in the presence of a high concentration of glucose (10 g L^-1^, Figure S7) and therefore the faster growth observed in CRISPRi strains compared to the control can be attributed to *crc* downregulation during glucose + *p*CA or FA conditions and not due to the presence of additional carbon source.

**Figure 3.**
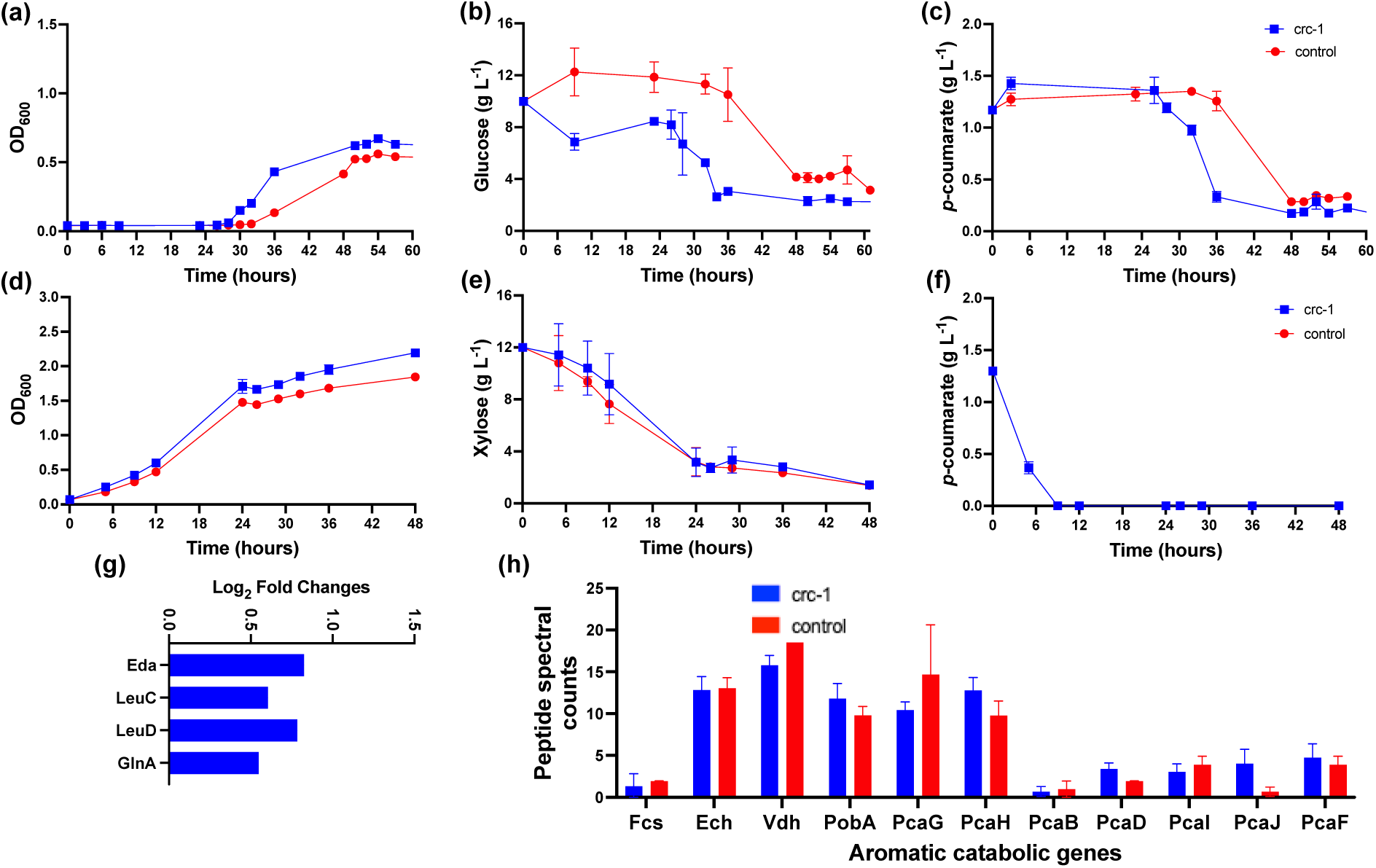
Growth performance (Phenotype) of *P. putida* M2 CRISPRi (crc-1) and wild-type strains showing OD_600_ and substrate consumption in glucose + *p*-coumarate (a, b, c) and xylose + *p*-coumarate (d, e, f) as carbon sources. The data represent the mean value of three replicates and error bars represent the standard deviation. The significantly upregulated (p ≤ 0.05, log_2_FC ≥ 0.5) proteins in the crc-1 strain compared to the control strain grown in glucose + *p*-coumarate (g). The log_2_FC was calculated from three biological replicates for each condition. The peptide spectral counts of aromatic catabolic genes in crc-1 and control strains when grown in glucose + *p*-coumarate (h).

The MS-based proteomics showed that Eda was significantly upregulated (0.8 log_2_FC, Figure 3g, Table S22) in crc-1 which might explain faster glucose consumption than the control strain. The proteins involved in amino acids biosynthesis (acetolactate synthase, glutamine synthetase, 3-isopropylmalate dehydratase, 3-isopropylmalate dehydrogenase, aspartate kinase) and transport were also significantly upregulated in the CRISPRi strains possibly explaining faster growth compared to the control (Table S22). However, there was no significant difference in the aromatic catabolic and the β-ketoadipate proteins between the CRISPRi strains (except upregulation of PcaD in the crc-2 strain) and the control grown in media containing glucose + *p*CA (Figure 3h).

### The abundance of sRNAs was proportional to the strength of carbon catabolite repression

Bacterial sRNAs are noncoding RNAs, ranging from 50-500 nt, that are found to be involved in gene regulation (26). In Pseudomonads, the Crc protein interacts with Hfq, RNA chaperon protein, creating a stable Hfq/Crc/mRNA complex (20). The strength of CCR is inversely proportional to the abundance of one or more sRNAs as they antagonize Crc by sequestering Hfq (22). Their abundance is low under conditions that elicit strong carbon catabolite repression and high in the absence of catabolic repression (27–29). Moreno et al (27) showed that the levels of sRNAs, CrcY and CrcZ, in *P. putida* PBA1 were low in cells growing exponentially in lysogeny broth (LB, strong CCR conditions) than in minimal media containing citrate or succinate (non-repression conditions). Under non-repressing conditions, CrcY and CrcZ sequestered Crc and impeded its binding to target mRNAs.

In the current study, the abundance of CrcY and CrcZ in KT2440 correlated with catabolite repression observed in the phenotypic study as well as with the proteomics results. The CrcY and CrcZ levels were lowest in the dual carbon condition compared to that in the single substrate condition (Figure 4), showing potential catabolite repression. A search of M2 sRNA sequences identified homologous sequences with 94% and 97% similarities to KT2440 CrcY and CrcZ, respectively. The expression of these homologues in M2 was lower under the glucose + *p*CA condition (strong CCR) than when only *p*CA or glucose (non-repressing condition) was present, possibly allowing the Crc protein to block the translation of the target mRNAs. On the other hand, when *p*CA or glucose was provided as the sole carbon source, the CrcY and CrcZ levels were higher, sequestering Crc by binding to it with high affinity as catabolite repression is not needed. Furthermore, CrcZ and CrcY in both KT2440 (six and four in CrcZ and CrcY, respectively) and M2 (six repeated motifs in both sRNA homologues) contain several repeated AANAANAA boxes, which have been described as Crc binding site (20, 30). These preliminary results suggest that the levels of sRNAs might have a role in controlling the amount of free Crc protein and, therefore, the strength of catabolite repression in KT2440 and M2 as already demonstrated in other *P. putida* strains.

**Figure 4.**
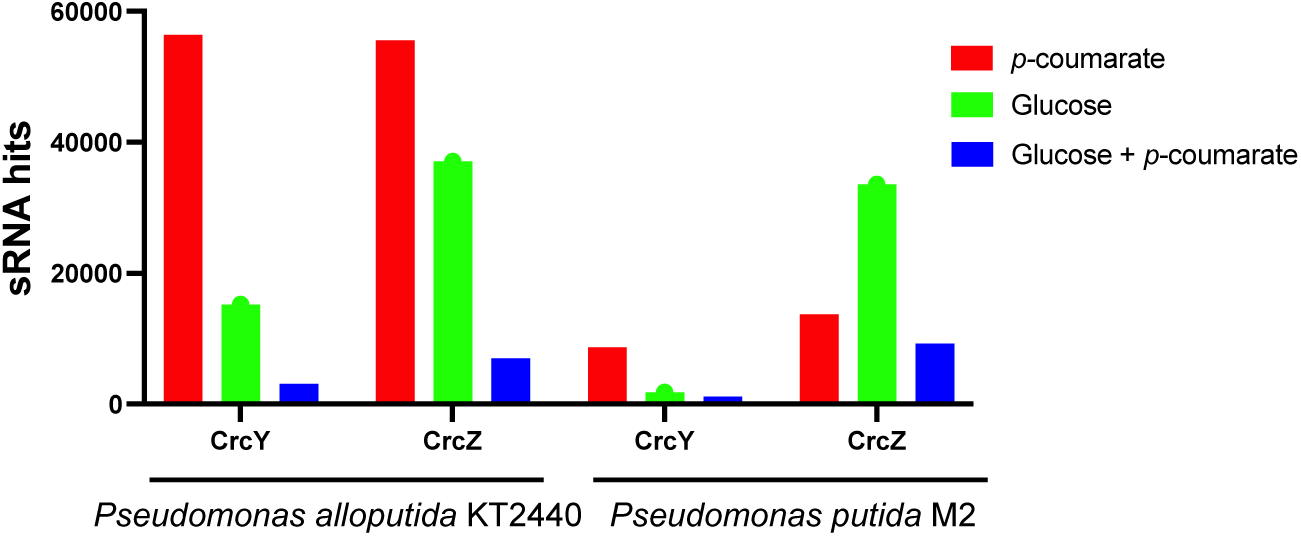
The abundance of sRNAs, CrcY and CrcZ, in *P. alloputida* KT2440 and *P. putida* M2 when grown in the presence of *p*-coumarate, glucose, or glucose + *p*-coumarate.

## 3. DISCUSSION

Pseudomonads grow and compete in diverse habitats due to their flexible metabolic pathway and have a global regulatory mechanism referred to as CCR that ensures that the most energy-efficient carbon source is utilized from a mixture of nutrient sources. This study demonstrated that M2 can co-metabolize sugars and aromatics; however, both sugars and aromatics were not completely consumed, except for aromatics in the presence of xylose. Unlike glucose-aromatic, the aromatics were consumed rapidly, and the aromatic intermediates accumulated transiently in the xylose-aromatic mixtures. The proteomics results also showed that glucose repressed all the aromatic catabolic and β-ketoadipate proteins while xylose inhibited only some of the aromatic catabolic proteins only. Therefore, glucose exerted a stronger CCR effect over the consumption of aromatics compared to xylose. The difference in CCR between glucose and xylose could be due to the difference in reducing power generated during their catabolism and their dependence on the aromatic catabolic pathways for their energy need. A multi-omics approach integrating proteomics, ^13^C-metabolomics, and quantitative flux analysis will provide a detailed understanding of carbon partitioning and the metabolic network during the co-utilization of sugars and aromatics in M2.

In the case of glucose and aromatics, there was a long lag phase after which consumption of both substrates commenced simultaneously and followed a similar trend indicating cross-catabolite repression. This mode of metabolism differs from the CCR where one carbon is strictly preferred over the other as observed in other cases (15, 31, 32). For example, *Pseudomonas putida* CSV86 preferentially consumes aromatics (naphthalene and benzyl alcohol) and organics acids over glucose where glucose metabolizing enzymes such as glucose-6-phosphate dehydrogenase and glucose transport protein were repressed during aromatics or organics acids utilization (31, 32). The phenomena of simultaneous CCR have been observed in KT2440 strains where glucose and toluene (33) and glucose and benzoate (34, 35) were simultaneously metabolized while repressing each other’s metabolism but in a manner that optimizes biomass growth. In KT2440 (pWW0), toluene-induced repression of glucose metabolism was directed to the glucokinase branch of glycolysis, while glucose inhibited TOL plasmid Pu promoter for toluene catabolism (33). A segregated metabolic flux was observed in KT2440-JD1 grown on glucose and benzoate where glucose-derived carbons were compartmentalized in the upper Embden-Meyerof-Parnas (EMP) and the pentose-phosphate (PP) pathways and the benzoate-derived carbons primarily populated the TCA cycle and the glyoxylate shunt (34). In our study with M2, the proteome analysis demonstrated that the presence of sugars repressed the translation of both the upper aromatic catabolic pathway and β-ketoadipate genes required for the aromatics assimilation including TCA and gluconeogenesis (in the case of xylose) proteins while the effect of aromatics was directed towards the Entner-Doudoroff (ED) branch (particularly the *ed*a gene) and the *xylX* gene, required for glucose and xylose metabolism, respectively, as well as the amino acid biosynthesis and catabolic pathways.

In M2, aromatic monomers such as *p*CA and VA are funneled into central intermediate PCA, which is further catabolized through the β-ketoadipate pathway to the TCA cycle to yield acetyl-CoA (10). The conversions of 4-HBA and VA to PCA catalyzed by PobA and VanAB, respectively, are considered the rate-limiting steps in KT2440 and have been identified as targets for Crc regulation in KT2440 (14). PobA and VanAB are the only enzymes in the aromatic catabolic pathway that require reducing equivalents for their activity, therefore they may be subject to tighter regulation by Crc when selecting the most energy-efficient carbon substrate. Replacing the native PobA in KT2440 with PraI from *Paenibacillus* sp. JJ-1b, which has a broader nicotinamide cofactor preference, relieved this bottleneck and improved muconate production from *p*CA and glucose (36). In M2, PobA and VanAB were also significantly downregulated, however, more work is needed to identify metabolic bottlenecks preventing the complete metabolism of aromatics in the presence of glucose in M2.

The global regulatory protein Crc plays a key role in CCR in Pseudomonads where Crc has been shown to inhibit the translation of mRNA encoding the catabolic enzymes, transcriptional regulators, as well as transporters required for substrates to enter the cell (37–39). The phenotypic characterization demonstrated that downregulating the *crc* gene led to faster growth and metabolism of CRISPRi strains compared to the control during the co-utilization of glucose and *p*CA or FA. The faster growth was further supported by evidence of upregulation of proteins involved in amino acids biosynthesis and transport while the upregulation of Eda potentially led to faster glucose consumption. However, the CRISPRi strains still exhibited an accumulation of aromatic intermediates and incomplete substrate consumption which suggests there might be additional yet unknown regulatory elements that control CCR. While alleviating CCR by deleting the *crc* gene has been shown to be beneficial to ensure efficient metabolism and improve product yield (14, 29, 40, 41), there are also contrary results that showed no positive effect after *crc* deletion (18, 42–44). Castillo et al.(33) showed that repression of glucose metabolism by toluene is mediated through the Crc protein while glucose-mediated repression of toluene degradation requires PtsN protein. Besides Crc, some potential regulators related to catabolite repression in *P. putida* are the cytochrome o ubiquinol oxidase (Cyo), the phosphoenolpyruvate: sugar phosphotransferase (Pts) system (44–48). Therefore, CCR regulation is a complex phenomenon that can be carbon source- and strain-specific and more often involves more than one regulatory element. The mechanism of Crc-mediated CCR and other potential regulatory mechanisms are still unknown in M2 and need to be further investigated in future research. Besides alleviating CCR, it is also necessary to broaden the substrate spectrum, enhance substrate uptake, and improve tolerance to toxic compounds that might be present in biomass hydrolysates or lignolysates to create an efficient and robust single strain-based biocatalyst for upgrading lignocellulosic biomass (49). Overexpressing gene encoding aromatic transporter (e.g. PcaK, the 4-HBA transporter (50)) was shown to improve the bioconversion of pCA to 2,4 pyridine dicarboxylic acid in engineered KT2440 (native *pcaHG* was replaced with *ligAB* from *Sphingobium* sp. SYK-6) and avoid 4-HBA accumulation (51). Future research should identify transporter genes and evaluate if overexpression or heterologous expression of transporter genes would improve bioconversion.

The difference in sRNA abundances between different carbon sources in both KT2440 and M2 followed the same trend, however, the difference was more distinct in KT2440 compared to that in M2. While we have no experimental evidence to explain these differences, however, there may be other potential sRNAs in M2 which played a greater role in CCR. Previous studies have used bioinformatics tools for sRNAs prediction (52, 53) and experimental approaches involving shotgun cloning approach combined with Northern blotting and RT-PCR (28, 54), to identify sRNAs involved in CCR in *Pseudomonas* strains and can provide directions for future studies in M2.

As demonstrated in this study, M2 may be used as a platform biocatalyst to produce industrially relevant products from lignocellulosic biomass due to its versatile sugar and aromatic metabolism. Our findings provide important insights into the physiology and metabolism of M2 during sugar and aromatic co-utilization. However, incomplete consumption and accumulation of aromatic intermediates due to CCR may hinder its industrial application. Simultaneous and complete utilization of sugar and aromatics could also be beneficial for bioproduct formation, one substrate could be used for energy for cell growth while the other substrate for product formation, a strategy already being used in KT2440 (14). As M2 is amenable to genetic manipulation via CRISPRi as shown by our study and previous publications (10, 25), this study provides a framework for use of metabolic engineering tools to alleviate CCR without causing other metabolism imbalances.

## 4. MATERIALS AND METHODS

### 4.1. Culturing conditions for *P. putida* M2 and *P. alloputida* KT2440 and growth measurements

All microbial cultivation experiments were conducted in triplicates at 30°C using 10 mL 0.2 μm filter sterilized modified M9 mineral medium (final pH adjusted to 7) with the following composition: (NH_4_)_2_SO_4_ (1.0 g L^-1^), KH_2_PO_4_ (1.5 g L^-1^), Na_2_HPO_4_ (3.54 g L^-1^), ammonium ferric citrate (0.06 g L^-1^), MgSO_4_·7H_2_O (0.2 g L^-1^), CaCl_2_·2H_2_O (0.01 g L^-1^), and trace elements H_3_BO_3_ (0.3 mg L^-1^), CoCl_2_·6H_2_O (0.2 mg L^-1^), ZnSO_4_·7H_2_O (0.1 mg L^-1^), MnCl_2_·4H_2_O (0.03 mg L^-1^), NaMoO_4_·2H_2_O(0.03 mg L^-1^), NiCl_2_6H_2_O (0.02 mg L^-1^), and CuSO_4_·5H_2_O (0.01 mg L^-1^). The M2 and KT2440 strains were inoculated from glycerol stocks onto an LB agar plate and a single colony from the plate was grown in 5 mL of LB medium (first preculture) overnight with continuous shaking at 200 rpm. The first preculture (100 μL) was inoculated into 5 mL freshly prepared M9 minimal medium with corresponding carbon source to prepare the second preculture and incubated overnight. Finally, the growth curve and growth kinetics experiments were started the following day from the second preculture at a desired starting optical density (0.025-0.05) in 50 mL glass tubes and 48 well plates with 250 uL cell culture in each well, respectively. The minimal media was supplemented with a mixture of sugar (glucose or xylose) and monomeric aromatics (*p*CA, 4-HBA, FA, or VA) as model lignocellulose-derived substrates at a concentration of ∼10 g L^-1^ (sugars) and ∼1 g L^-1^ (aromatics). For the growth curve experiments, the cell culture samples were taken at different time points to determine growth spectrophotometrically (SpectraMax M2, Molecular Devices, San Jose, CA) by measuring absorbance at 600 nm (OD_600_). Growth kinetics was done using a Synergy plate reader (BioTek Instruments, Winooski, VT, USA).

Quantification of sugars was performed using high performance liquid chromatography (HPLC) using Agilent 1260 Infinity system (Santa Clara, CA, USA) equipped with an Aminex HPX-87H (300 mm × 7.8 mm) column (Bio-Rad, Hercules, CA) and a refractive index detector heated at 35 °C. An aqueous solution of 4 mM sulfuric acid was used as the mobile phase (0.4 mL min^−1^, column temperature of 25 °C). The amount of monomeric aromatic compounds was analyzed using Agilent 1200 system with Eclipse Plus Phenyl-Hexyl (250 mm length, 4.6 mm diameter, 5 µm particle size) column kept at 50°C. The mobile phase consisted of binary system phases with 10 mM ammonium acetate in 0.07% formic acid (Solvent A) and 10 mM ammonium acetate in 90% acetonitrile and 0.07% formic acid (Solvent B). The mobile phase gradient method was as follows: 30% B (0 min; 0.5 mL min^−1^), 80% B (12 min; 0.5 mL min^−1^), 100% B (12.1 min; 0.5 mL min^−1^), 100% B (12.6 min; 1 mL min^−1^), 30% B (12.8 min; 1 mL min^−1^), 30% B (15.6 min; 1 mL min^−1^). The metabolite concentration was quantified from peak areas using calibration curves generated by each metabolite standard.

### 4.2. Shotgun proteomics analysis using LC-MS

All strains were grown in three biological replicates at 30°C following a similar procedure as stated in Section 4.1 starting with an LB agar plate followed by the first and second pre-culture before growing them in 10 mL M9 minimal media supplemented with corresponding carbon sources (Table S1) at 30°C. The cells were harvested by centrifugation at OD between 0.4-0.5 and stored at −80°C until sample preparation. Proteins from the samples were extracted using a previously described chloroform/methanol precipitation method (55). Extracted proteins were resuspended in 100 mM ammonium bicarbonate buffer supplemented with 20% methanol, and protein concentration was determined by the DC assay (BioRad). Protein reduction was accomplished using 5 mM tris 2-(carboxyethyl)phosphine (TCEP) for 30 min at room temperature, and alkylation was performed with 10 mM iodoacetamide (IAM; final concentration) for 30 min at room temperature in the dark. Overnight digestion with trypsin was accomplished with a 1:50 trypsin:total protein ratio. The resulting peptide samples were analyzed on an Agilent 1290 UHPLC system coupled to a Thermo scientific Obitrap Exploris 480 mass spectrometer for discovery proteomics (56). Briefly, 20 µg of tryptic peptides were loaded onto an Ascentis® (Sigma–Aldrich) ES-C18 column (2.1 mm × 100 mm, 2.7 μm particle size, operated at 60°C) and were eluted from the column by using a 10 min gradient from 98% buffer A (0.1 % FA in H2O) and 2% buffer B (0.1% FA in acetonitrile) to 65% buffer A and 35% buffer B. The eluting peptides were introduced to the mass spectrometer operating in positive-ion mode. Full MS survey scans were acquired in the range of 300-1200 m/z at 60,000 resolution. The automatic gain control (AGC) target was set at 3E06 and the maximum injection time was set to 60 ms. Top 10 multiply charged precursor ions (2–5) were isolated for higher-energy collisional dissociation (HCD) MS/MS using a 1.6 m/z isolation window and were accumulated until they either reached an AGC target value of 1e5 or a maximum injection time of 50 ms. MS/MS data were generated with a normalized collision energy (NCE) of 30, at a resolution of 15,000. Upon fragmentation precursor ions were dynamically excluded for 10 s after the first fragmentation event. The acquired LCMS raw data were converted to mgf files and searched against the latest uniprot P. putida KT2440 protein database or JGI P. putida M2 protein database (IMG submission ID: 236130) with Mascot search engine version 2.3.02 (Matrix Science). The resulting search results were filtered and analyzed by Scaffold v 5.0 (Proteome Software Inc.). The normalized spectra count of identified proteins were exported for relative quantitative analysis. The proteomics data were analyzed using a customized Python script to calculate log_2_ FC values and identify proteins that were significantly downregulated in the dual substrate conditions compared to the single substrate condition. Only proteins with spectral counts above 5 were retained in the analysis. Proteins were identified to be significantly differentially expressed when they displayed a log_2_ FC ≥ 0.5 across at least two out of three biological replicates with the p-value cut-off of 0.05.

### 4.3. CRISPR interference-based *crc* gene repression

*Escherichia coli* TOP10 cells were used as the primary host for gene cloning and plasmid isolation. Kanamycin (50 μg ml^-1^) was used for the selection of plasmid-harboring cells. *E. coli* competent cell preparation and transformation were performed using standard molecular biology techniques (57). Electroporation (1 mm gap cuvette, with parameters set at the resistance of 200Ω, Voltage 1.8 KV, capacitance of 25 μFD for 4.0-5.0 ms) was used for the transformation of competent M2 cells with selected plasmids. Cells were incubated in LB medium for 1.5 h at 30°C and 200 rpm on a shaker incubator and plated on LB plates with kanamycin for selection. Q5 High-Fidelity 2X and OneTaq^TM^ 2X Master Mix (New England Biolabs; Ipswich, MA, USA) were used according to the manufacturer’s instructions for PCR amplification during gene cloning and colony PCR to confirm gene cloning, respectively. NEBuilder® HiFi DNA Assembly Master Mix (New England Biolabs) was used to assemble plasmids. QIAprep Spin miniprep kit and QIAquick gel extraction kit (Qiagen, Hilden, Germany) were used according to the manufacturer’s protocol to isolate plasmids and to separate PCR products from agarose gels, respectively. The sequences of all plasmids and DNA constructs were confirmed by Sanger sequencing (Genewiz, CA, USA).

The CRISPR interference system utilizing the type II dCas9 homologs from *Streptococcus pyogenes* (pRGPspdCas9bad, Table S23) previously developed for KT2440 and M2 (24, 25) was used as the backbone plasmid to develop a dual inducible CRISPR/Cas9 interference-based gene repression system in M2. Expression of spdCas9 was under the control of the P_tac_ (Isopropyl β-D-1-thiogalactopyranoside (IPTG)-inducible) promoter while sgRNA expression was under the control of the P*_araC/bad_* (arabinose-inducible) promoter. The wild-type M2 strain carrying an empty pRGPspdCas9bad plasmid served as the control. An open reading frame (Ga0436255_01_2263068_2263847) was found which had 95% sequence similarity to the KT2440 *crc* sequence in the M2 genome. Three sgRNA (20 bp) targeting three different regions of the *crc* gene (Figure S8), resulting in three recombinant strains: crc-1, crc-2, and crc-3 were designed and cloned into the plasmid by PCR amplifications using primers listed in Table S24. One sgRNA targeted a region close to the start codon at the 5’ end (crc-1) and two inside the ORF (crc-2 and crc-3) of the *crc* gene. The constructed plasmids were electroporated into M2 to generate the CRISPRi strains. The recombinant CRISPRi strains were grown in minimal media supplemented with corresponding carbon sources (glucose + *p*CA, glucose + FA, and xylose + *p*CA), and both inducers (∼3.5 g L^-1^ arabinose and 1 mM IPTG) for phenotypic characterization (following a similar protocol as stated in Section 4.1) and proteome comparison (glucose, *p*CA, or glucose + *p*CA as carbon sources) with the control strain.

The oligonucleotides were ordered from Integrated DNA Technologies (IDT, San Diego, CA, USA). All the bacterial strains, plasmids, and primers used in this study can be found in Tables S23 and S24 in the Supplementary information.

### 4.4. small RNA sequencing

M2 and KT2440 cells were grown in minimal media containing either glucose, *p*CA, or glucose + *p*CA, and cell pellets were submitted to CD genomics (NY, USA) for total RNA extraction and sRNA sequencing. Briefly, total RNA was isolated using Qiagen miRNeasy mini kit (Hilden, Germany) following the manufacturer’s protocol, quantified by NanoDrop^TM^ 2000 (Thermo Fisher Scientific, Waltham, MA, USA), and quality assessed with agarose gel electrophoresis. The preparation of the sRNA-enriched library was done with NEBNext Ultra II RNA kit (New England Biolabs, Ipswich, MA, USA) following the manufacturer’s recommendation which included cDNA synthesis, 3’ and 5’ adapter ligation, PCR enrichment of adapter-ligated RNA, and size fractionation (80-400 bp) with Pippin Prep to specifically enrich for sRNAs. The library quantity was measured by KAPA SYBR FAST qPCR while the quality was evaluated by TapeStation D1000 ScreenTape (Agilent Technologies, CA, USA). Equimolar pooling of libraries was performed based on the QC values and sequenced on Illumina NovaSeq 6000 (Illumina, CA, USA) with a read length configuration of 150 paired end for 10 million paired-end reads per sample.

Illumina adapter sequences and small RNA library adapter sequences were removed from raw sequencing reads using Fastp and Cutadapt, respectively. The coding mRNA sequences were removed by aligning the quality-filtered RNA-seq reads with the reference genome. First, we ran BLASTX against sequences similar to CrcY and CrcZ, the sRNAs found in KT2440 (27), and determined their abundances in both M2 and KT2440 under different carbon conditions.

## Data availability

The generated mass spectrometry proteomics data have been deposited to the ProteomeXchange Consortium via the PRIDE (58) partner repository with the dataset identifier PXD041554 and 10.6019/PXD041554. The recombinant strains used in this study have been deposited in the public instance of the JBEI registry (http://public-registry.jbei.org/). The raw small RNA sequences are available in the NCBI short read archive under accession number PRJNA953382.

## Acknowledgments

This work was as part of the Department of Energy (DOE) Joint BioEnergy Institute (http://www.jbei.org) supported by the U.S. DOE, Office of Science, Office of Biological and Environmental Research, through Contract No. DE-AC02-05CH11231 between Lawrence Berkeley National Laboratory and the U.S. DOE. The U.S. Government retains and the publisher, by accepting the article for publication, acknowledges that the U.S. Government retains a nonexclusive, paid-up, irrevocable, worldwide license to publish or reproduce the published form of this manuscript or allow others to do so, for the U.S. Government purpose.

